# Isw2 and Ino80 chromatin remodeling factors regulate chromatin, replication, and copy number at the yeast ribosomal DNA locus

**DOI:** 10.1101/291971

**Authors:** Sam Cutler, Laura J Lee, Toshio Tsukiyama

## Abstract

In the budding yeast *Saccharomyces cerevisiae*, ribosomal RNA genes are encoded in a highly repetitive tandem array referred to as the ribosomal DNA (rDNA) locus. The yeast rDNA is the site of a diverse set of DNA-dependent processes, including transcription of ribosomal RNAs by RNA Polymerases I and III, transcription of non-coding RNAs by RNA Polymerase II, DNA replication initiation, replication fork blocking, and recombination-mediated regulation of rDNA repeat copy number. All of this takes place in the context of chromatin, but relatively little is known about the roles played by ATP-dependent chromatin remodeling factors at the yeast rDNA. In this work, we report that the Isw2 and Ino80 chromatin remodeling factors are targeted to this highly repetitive locus. We characterize for the first time their function in modifying local chromatin structure, finding that loss of these factors affects the occupancy of nucleosomes in the 35S ribosomal RNA gene and the positioning of nucleosomes flanking the ribosomal origin of replication. In addition, we report that Isw2 and Ino80 promote efficient firing of the ribosomal origin of replication and facilitate the regulated increase of rDNA repeat copy number. This work significantly expands our understanding of the importance of ATP-dependent chromatin remodeling for rDNA biology.

**Author Summary:** To satisfy high cellular demand for ribosomes, genomes contain many copies of the genes encoding the RNA components of ribosomes. In the budding yeast *Saccharomyces cerevisiae*, these ribosomal RNA genes are located in the “ribosomal DNA locus”, a highly repetitive array that contains approximately 150 copies of the same unit, in contrast to the single copies that suffice for most genes. This repetitive quality creates unique regulatory needs. Chromatin structure, the packaging and organization of DNA, is a critical determinant of DNA-dependent processes throughout the genome. ATP-dependent chromatin remodeling factors are important regulators of chromatin structure, and yet relatively little is known about how members of this class of protein affect DNA organization or behavior at the rDNA. In this work, we show that the Isw2 and Ino80 chromatin remodeling factors regulate two features of chromatin structure at the rDNA, the occupancy and the positioning of nucleosomes. In addition, we find that these factors regulate two critical processes that function uniquely at this locus: DNA replication originating from within the rDNA array, and the regulated increase of rDNA repeat copy number.

## Introduction

In exponentially growing cells, the enormous cellular demand for ribosomes is reflected in the proportion of resources dedicated to their production. For example, the production of ribosomal RNAs (rRNAs) accounts for an estimated 60% of all transcriptional activity in cycling yeast cells [1]. Because single genomic copies of rRNA genes would not support such large volumes of transcriptional output, eukaryotic genomes have evolved to include highly repetitive clusters of rRNA genes, termed the ribosomal DNA (rDNA) locus. In a typical cell of the budding yeast *Saccharomyces cerevisiae*, the rDNA locus comprises approximately 150-200 tandem repeats (Fig 1A). Each repeat contains a 35S ribosomal RNA (rRNA) gene, transcribed by RNA Polymerase I (Pol I), and an inter-genic spacer (IGS), split into IGS1 and IGS2 regions by the 5S rRNA gene, which is transcribed by RNA Polymerase III (Pol III). Due to its large size and repetitive nature, the rDNA locus has unique regulatory needs, and the IGS1 and IGS2 regions contain genetic elements that are critical to addressing these needs.

**Fig 1.**
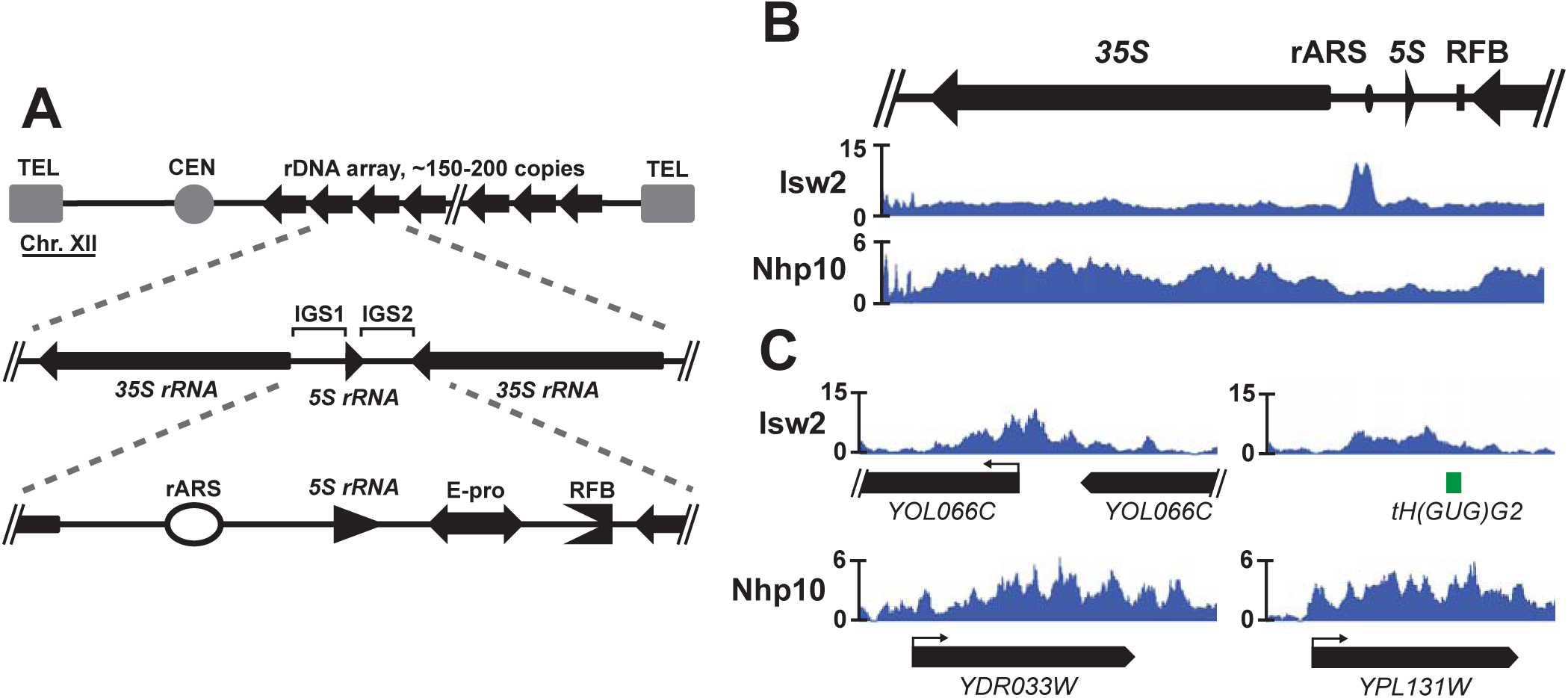
The Isw2 and Ino80 chromatin remodeling complexes are targeted to the rDNA locus. (A) A schematic drawing of the rDNA locus in *S. cerevisiae*. In a typical yeast cell, the rDNA accounts for approximately 1.5 Mb of chromosome XII, comprised of a tandem array of ~150 copies of the rDNA repeat. Each repeat contains a 35S rRNA gene and an inter-genic spacer (IGS) region in between adjacent 35S genes, itself split into IGS1 and IGS2 regions by the 5S rRNA gene. IGS1 contains the ribosomal origin of replication, or autonomously replicating sequence (rARS), and IGS2 contains the bi-directional RNA Polymerase II promoter, E-pro, and a replication fork block (RFB). (B) The Isw2 subunit of the Isw2 complex and the Nhp10 subunit of the Ino80 complex were each FLAG-tagged, chromatin immuno-precipitated, and deep-sequenced (ChIP-seq). (C) Representative ChIP-seq signals of Isw2 and Nhp10 at single copy targets outside of the rDNA.

Without an origin of replication (autonomously replicating sequence, or ARS), replicating the rDNA array would require replication forks to traverse multiple megabases of DNA from either end of the array. To avoid this, IGS1 contains a ribosomal ARS (rARS). As a consequence, the approximately 150 ARSs in a typical rDNA array account for nearly one third of all genomic origins of replication. Of these rARSs, only around 20% will fire in any given round of cell division [2, 3]. Because replication factors are limiting during each S-phase [4], firing of too many rARSs would take vital replicative resources away from other parts of the genome, raising the risk of delayed or incomplete replication. If too few rARSs fire, replication of the rDNA array may be delayed or incomplete [5]. Thus, properly striking this balance by regulating origin efficiency at the rDNA has critical consequences for global genome stability.

Genome stability is also affected by the size of the rDNA array [6]. Because of this, a mechanism exists to change the size of the array by adding or removing copies of the rDNA repeat if needed. The IGS2 region contains two genetic elements that are critical for this process: a bi-directional RNA Pol II promoter, E-pro, and a replication fork block (RFB). The RFB pauses replication forks moving through the IGS toward the 3’ end of the 35S gene, but allows forks coming from within the adjacent 35S gene, and thus moving in the same direction as 35S transcription, to pass through or merge with paused forks. This activity is thought to prevent head-on collisions between replication machinery and densely loaded Pol I machinery in the highly transcribed 35S [3, 7]. In addition, forks paused at the RFB are the sites of targeted DNA double-strand breaks (DSBs). The level of transcription from the adjacent E-pro promoter influences the mechanism by which these targeted DSBs are repaired, which in turn influences whether a repeat is added to or removed from the rDNA array, or whether the copy number remains unchanged [8].

All DNA-dependent processes occurring at the rDNA, including transcription by multiple RNA polymerases, origin firing, and changes in rDNA copy number, happen in the context of chromatin structure. The Sir2 and Rpd3 histone deacetylases (HDACs) have well-established roles in regulating rDNA chromatin structure, origin activity, and copy number maintenance [5, 8–10]. In addition, rDNA biology is regulated by ATP-dependent chromatin remodeling factors, which use the energy of ATP hydrolysis to modify the position and histone composition of nucleosomes. In humans, the nucleolar remodeling complex (NoRC) positions nucleosomes and recruits histone methyltransferase and histone deacetylase activity to promote rDNA silencing [11, 12]. In yeast, the SWI/SNF [13], Isw1, Isw2, and Chd1 [14] complexes have been implicated in regulating transcription of rRNAs. Until now, no remodeling factors have been shown to modify chromatin structure at the yeast rDNA or to affect any DNA-dependent processes beyond rRNA transcription at this locus.

In this work, we show that the Isw2 and Ino80 ATP-dependent chromatin remodeling factors regulate chromatin structure at the rDNA. The Isw2 complex is known to slide nucleosomes over gene promoters [15], an activity that generally represses transcription, both for coding genes [16, 17] and antisense transcripts [18]. The Ino80 complex slides and evicts nucleosomes and removes the histone variant, H2A.Z [19–22]. Ino80 is also involved in regulating the checkpoint response following DNA damage, DNA damage repair, and DNA replication [23–25]. Isw2 and Ino80 function together to promote replication of late-replicating regions of the genome in the presence of replication stress and to attenuate the S-phase checkpoint response [26, 27]. Here, we show that both Isw2 and Ino80 are targeted to the ribosomal DNA locus in distinct patterns, primarily characterized by striking enrichment of Isw2 around the rARS and of Ino80 through the 35S gene. Further, we report for the first time that these remodeling factors affect local chromatin structure, as loss of the factors increases nucleosome occupancy in the 35S and alters the positioning of nucleosomes flanking the rARS. We find that loss of Isw2 and Ino80 reduces the proportion of active rDNA repeats without affecting overall transcription of rRNAs, but that Isw2 and Ino80 positively contribute both to the efficiency of the rARS and to the rate of rDNA repeat copy number change. In sum, this study significantly expands our understanding of how ATP-dependent chromatin remodeling factors affect both chromatin structure and essential biological processes at the ribosomal DNA locus.

## Results

### The Isw2 and Ino80 chromatin remodeling complexes are targeted to the ribosomal DNA locus

All of the DNA-dependent processes that take place at the rDNA locus occur in the context of chromatin. Although HDACs such as Rpd3 and Sir2 have well-characterized functions in regulating chromatin structure, transcription, and copy number maintenance at the *S. cerevisiae* rDNA [8–10, 28], comparatively little is known about the roles played by ATP-dependent chromatin remodeling factors at this vitally important genomic locus. To address this, we performed chromatin immuno-precipitation followed by deep sequencing (ChIP-seq) to map where the Isw2 and Ino80 chromatin remodeling factors are targeted at the rDNA. We found that the namesake, catalytic subunit of Isw2 and Nhp10, a subunit wholly unique to the Ino80 complex [23], were both targeted to the rDNA (Fig 1B). The ChIP-seq signal for Isw2 was slightly above the genome-average throughout the 35S gene body. The pattern of targeting in the IGS included small peaks flanking the 5S gene and the region containing E-pro and RFB, but the most prominent signal was a striking, bimodal peak on top of and to one side of the rARS. Nhp10 was also present throughout the 35S gene body and showed a small peak around the 5S. Each protein’s ChIP-seq pattern at the rDNA was consistent with peaks elsewhere in the genome with regard to both shape and magnitude: Isw2 tended to have fairly defined peaks that rise well above the genome average, located in intergenic regions, and Nhp10 peaks were generally less prominent relative to the genome average and more diffusely spread throughout a transcription unit (Fig 1C). Given these distinct targeting patterns, we hypothesized that these ATP-dependent chromatin remodeling factors might have previously unknown functions at this highly repetitive, unique genomic locus.

### Isw2 and Ino80 affect nucleosome occupancy over the 35S rRNA gene

In light of the established functions of the Isw2 and Ino80 complexes, we first asked whether these chromatin remodeling factors affect nucleosome occupancy within the rDNA locus, as this feature of chromatin structure has well-established importance at the rDNA. Individual rDNA repeats canonically exist in one of two discrete states, being either highly occupied with nucleosomes and transcriptionally inactive, or heavily depleted of nucleosomes and highly transcriptionally active [29–31]. We assessed how nucleosome occupancy at the rDNA is affected by these two chromatin remodeling factors with ChIP-seq of histone H3 in wild-type, *isw2∆*, *nhp10∆*, and *isw2∆ nhp10∆* strains. This analysis revealed that nucleosome occupancy throughout the 35S gene body is appreciably increased in the *isw2∆ nhp10∆* double mutant compared to wild-type and single deletion strains (Fig 2A, left panel). Notably, this is the part of the rDNA in which the ChIP-seq signals of both chromatin remodeling factors most significantly overlap, suggesting the possibility that these factors may work together in this region.

**Fig 2.**
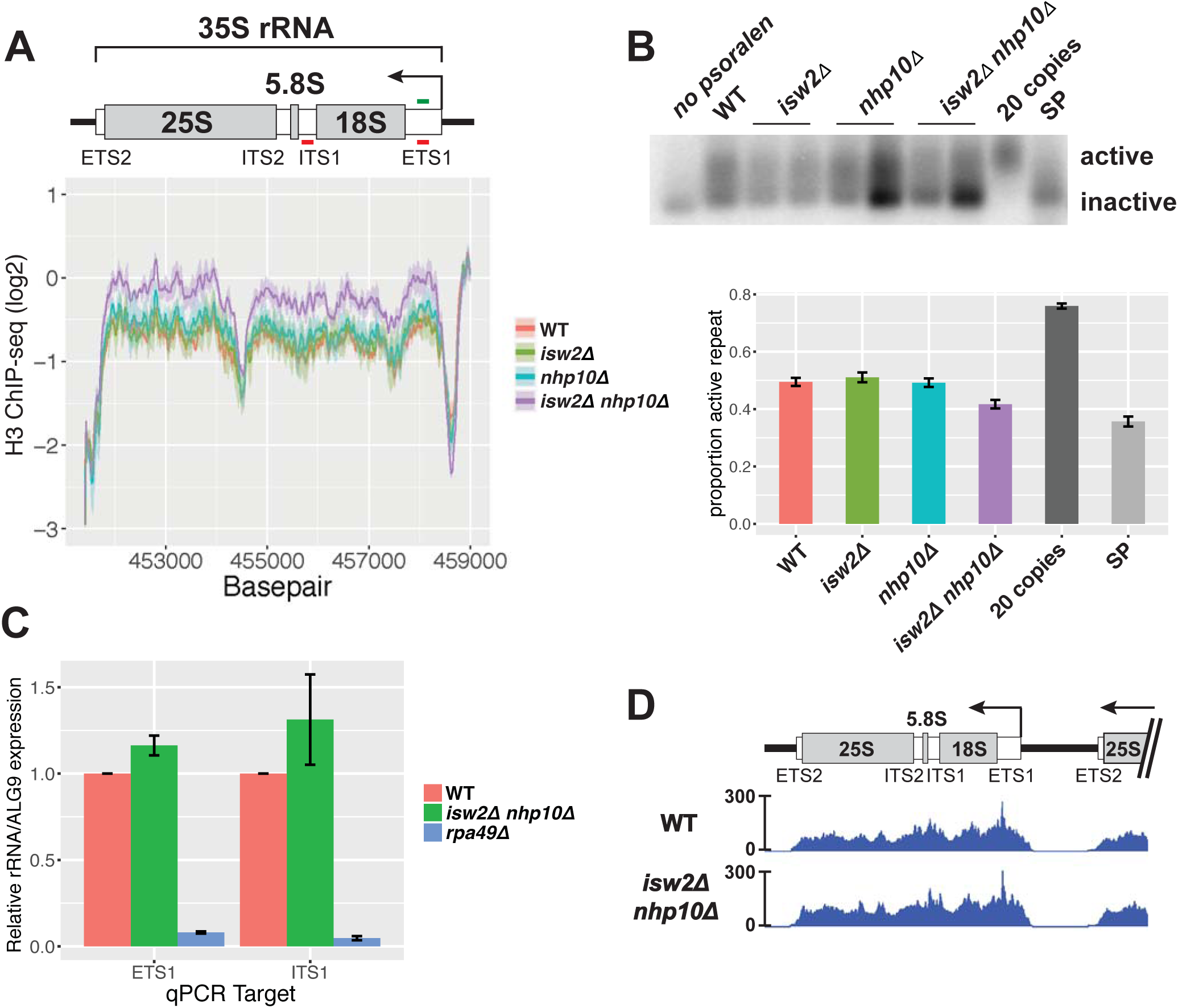
Nucleosome occupancy, but not transcription, is affected at the 35S rDNA in *isw2∆* and *nhp10∆* mutants. (A) Histone H3 ChIP-seq through the 35S rRNA gene. Line represents average log2 ChIP-seq signal at each base pair for two independent experiments, and the ribbon represents the standard error of the mean at each base pair. Schematic drawing of the 35S includes transcribed spacers that are removed during processing, as well as the mature 18S, 5.8S, and 25S rRNAs that are parts of complete ribosomes. ETS1 and ITS1 qPCR primer sets are indicated with red lines, and ETS1 hybridization probe, used in the Southern blot shown in 2B, indicated in green. In this and all following figures, “wild-type” has been abbreviated as “WT”. (B) Psoralen cross-linked DNA, digested with EcoRI and hybridized with a probe to the ETS1 region. Psoralen incorporates more readily into nucleosome-occupied, actively transcribed rDNA repeats, causing these bands to migrate more slowly than nucleosome-depleted, inactive repeats. Two independent isolates of each remodeling factor mutant are shown. For quantification, mean intensity of each band was measured with ImageJ software. Values for each genotype reflect between 3 and 5 biological replicates, and error bars represent standard error of the mean. (C) RT-qPCR measuring the ETS1 and ITS1 of the 35S pre-rRNA. (D) RNA Pol I ChIP-seq.

Given that rDNA repeats canonically exist in one of two discrete states that are associated with nucleosome occupancy, we hypothesized that the increased nucleosome occupancy in *isw2∆ nhp10∆* cells reflects a reduced ratio of active to inactive rDNA repeats. To test this, we used psoralen cross-linking, a well-established method in which DNA is treated with the DNA-intercalating compound, psoralen [29, 32]. Occupancy of chromatin by nucleosomes blocks incorporation of psoralen. Therefore, actively transcribed, nucleosome-depleted rDNA repeats become more heavily cross-linked with psoralen than inactive, nucleosome-occupied repeats. After digestion with appropriate restriction enzymes, Southern blotting, and hybridization with a probe targeting a region of the 35S gene unit, two discrete bands representing active and inactive repeats can be resolved [29, 32]. By this method, we found that *isw2∆ nhp10∆* cells have a reduced proportion of active repeats compared to wild-type, *isw2∆*, or *nhp10∆* cells (Fig 2B), consistent with the observed increase in H3 occupancy in double mutant cells. Based on these results, we concluded that the Isw2 and Ino80 chromatin remodeling factors increase the ratio of active to inactive rDNA repeats.

### Transcription of 35S ribosomal RNA is not affected by loss of Isw2 or Nhp10

Based on the reduced proportion of nucleosome-depleted rDNA repeats in the *isw2∆ nhp10∆* mutant, we hypothesized that these cells would also show reduced levels of 35S rRNA transcription. The 35S is transcribed as a single long transcript before being cleaved and folded in a series of processing steps to yield mature 18S, 5.8S, and 25S RNAs [33]. Because mature rRNAs are components of ribosomes and thus highly stable and abundant, nascent RNA needs to be measured to assess the transcription rate of rRNAs. The External Transcribed Spacer 1 (ETS1) and Internal Transcribed Spacer 1 (ITS1) sections of the 35S gene are transcribed but removed at early stages of rRNA processing. Levels of these RNA sequences thus reflect levels of nascent rRNA and are commonly used to measure the rate of 35S transcription [34, 35]. Adopting this approach, we performed reverse-transcription quantitative PCR (RT-qPCR) targeting parts of the ETS1 and ITS1 regions of the 35S pre-rRNA (Fig 2A). As expected, we found significantly reduced levels of both ETS1 and ITS1 in an *rpa49* deletion mutant, a strain known to have a reduced rate of RNA Pol I transcription [35, 36]. To our surprise, we did not see evidence of a significant difference in rates of 35S transcription in *isw2∆ nhp10∆* compared to wild-type (Fig 2C). To confirm this unexpected result by an independent method, we next performed ChIP-seq analysis of the Pol I subunit RPA190, and observed virtually identical profiles in *isw2∆ nhp10∆* and wild-type strains, with regard to both shape and overall levels (Fig 2D). Based on these results, we concluded that *isw2∆ nhp10∆* cells exhibit no significant defects in the rate of 35S transcription despite the observed differences in nucleosome occupancy and the proportion of nucleosome-occupied rDNA repeats in these mutants.

### Isw2 and Ino80 affect nucleosome positioning in the rDNA inter-genic spacer

In addition to nucleosome occupancy, nucleosome positioning is known to be affected by both of these chromatin remodeling factors [15, 20]. Therefore, we assessed nucleosome positioning at the rDNA by micrococcal nuclease (MNase) digestion followed by deep sequencing (MNase-seq). We interpret each size-selected, paired-end read as coming from a nucleosome-protected fragment of DNA, and so from each paired-end read, the nucleosomal dyad center was inferred and plotted, resulting in the profiles shown (Fig 3A). By this method, nucleosome positions appear strongly shifted at known Isw2 targets in *isw2∆* and *isw2∆ nhp10∆* mutants (S1 Fig). In contrast, no gross differences in nucleosome positions are observed throughout the 35S gene body (S2A Fig) or in the rDNA inter-genic spacer region (Fig 3A). Within the highly repetitive rDNA, sequencing data must be interpreted carefully, however, as it represents an average of the signal at all ~150 rDNA repeats in all cells sampled, and nucleosomes in only a fraction of those repeats may change positions in any given cell.

**Fig 3.**
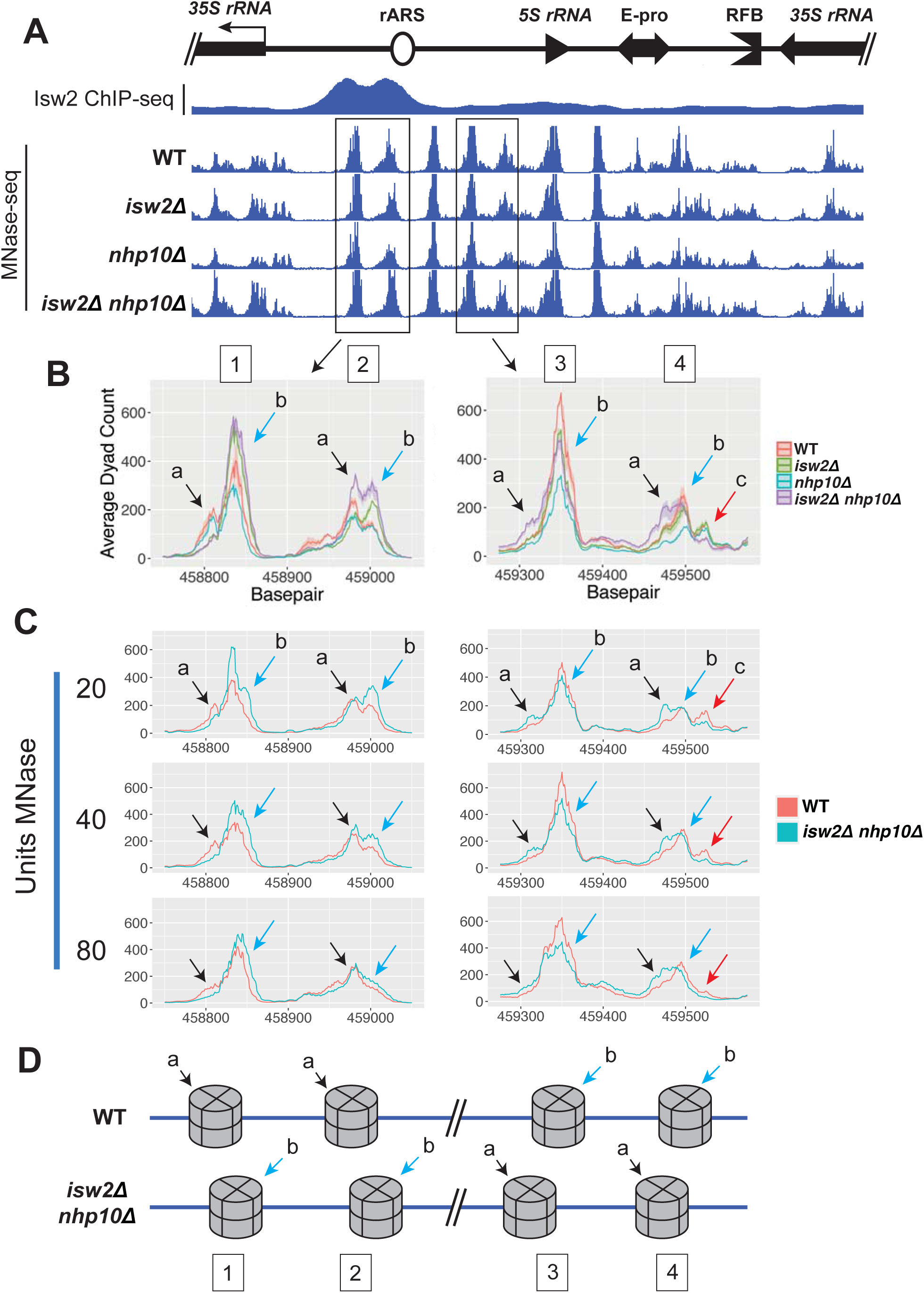
Isw2 and Ino80 affect nucleosome positioning in the rDNA inter-genic spacer. (A) Micrococcal nuclease digestion followed by deep-sequencing (MNase-seq) profiles in the IGS, with Isw2 ChIP-seq data overlaid. From each paired end sequencing read, the nucleosome dyad was inferred and plotted. (B) Ribbon plots, generated as described in Fig 2A, focused on two pairs of nucleosomes, indicated with boxes in Fig 3A. Each of the four tested strains has a characteristic profile of positioning at each of these four nucleosomes. Different sub-species of nucleosome positions are indicated with colored arrows and letters. (C) MNase-seq comparing wild-type and *isw2∆ nhp10∆* cells across three different strengths of MNase digestion. (D) Cartoon depicting different nucleosomal sub-species, highlighting the most striking differences in sub-species profiles between wild-type and *isw2∆ nhp10∆* cells.

To refine our analysis, we compared MNase-seq profiles for the tested strains using ribbon plots in which the primary line shows the average signal at each base pair across multiple biological replicates, and the ribbon represents the standard error of the mean for those replicates (Fig 3B). This method revealed striking differences in nucleosome positioning at the rDNA for two pairs of nucleosomes. One pair is in between the 35S promoter and the rARS, with each nucleosome substantially overlapping one of the two sub-peaks of the highly prominent Isw2 peak (Fig 3B, left panel, identified as nucleosomes 1 and 2). The other pair of affected nucleosomes is in the region between the rARS and the 5S gene, overlapping half of the short Isw2 peak encompassing the 5S (Fig 3B, right panel, nucleosomes 3 and 4). Each of these four MNase-seq dyad peaks appears to have two sub-species of nucleosome positions. We interpret each of these distinct sub-species as representing one of two distinct positions occupied by that nucleosome in different individual rDNA repeats in the array. Each of the four genotypes tested has a characteristic pattern of the relative heights of these two sub-species, which we propose reflects different proportions of rDNA arrays containing nucleosomes at either position.

At nucleosome 1, *isw2∆* and *isw2∆ nhp10∆* cells virtually only have sub-species 1*b*, while both wild-type and *nhp10∆* cells also have a significant signal for sub-species 1*a*. At nucleosome 2, wild-type cells predominantly have sub-species 2*a*, whereas *isw2∆* cells have more prominent signals for 2*b*, and *nhp10∆* and *isw2∆ nhp10∆* cells have roughly similar ratios of each sub-species. Nucleosome 3 resembles nucleosome 1, in that for some strains – in this case, wild-type, *isw2∆*, and *nhp10∆* cells – there is essentially only one sub-species, 3*b*, whereas only *isw2∆ nhp10∆* cells have a small but distinct sub-species 3*a*. For nucleosome 4, wild-type and *isw2∆* cells are very similar, with 4*b* dominating and 4*a* and 4*c* of similar, lower prominence, while *nhp10∆* cells have similar levels of 4c but proportionally reduced 4*a* and 4*b* peaks. Again, the double mutant is the most different among the tested strains, as 4*c* is barely detectable, while 4*a* is on par with 4*b* in *isw2∆ nhp10∆* cells. In sum, the overall trend among these mutants is that in *isw2∆ nhp10∆* cells, any given rDNA repeat is more likely to have nucleosomes positioned such that they are encroaching on the rARS. In contrast, in both *nhp10∆* and wild-type cells, these same nucleosomes are more likely to be positioned farther away from the rARS, and in *isw2∆* cells these nucleosomes have profiles somewhere in between wild-type and the double mutant.

Nucleosomes 3 and 4 are located in between the rARS and the 5S rRNA gene. Thus, the positioning shifts of these nucleosomes relative to the rARS also happen, in the opposite direction, relative to the 5S. In addition, there were slight strain specific differences at the nucleosome partially overlapping the 5S: sub-species *a* is higher than *b* in wild-type, *isw2∆*, and *nhp10∆* cells, but the sub-species are roughly equivalent in *isw2∆ nhp10∆* cells (S2B Fig). These differences in nucleosome positioning suggested that Isw2 and Ino80 might alter nucleosomes to regulate 5S transcription. Because this gene is only 120 bp in length and undergoes only minor processing before incorporation into ribosomes, it is difficult to distinguish between mature and nascent 5S rRNA transcripts. Therefore, to assess 5S transcription, we performed RNA Pol III ChIP-seq. Similarly to RNA Pol I levels at the 35S, Pol III levels at the 5S did not differ between wild-type and *isw2∆ nhp10∆* cells (S2C Fig). Thus, we conclude that Isw2 and Ino80 do not significantly affect transcription of ribosomal RNAs despite changes in chromatin structure around the transcription units.

It has been shown that the strength of MNase digestion can affect nucleosome mapping results, especially for nucleosomes that are highly MNase sensitive [37]. Because the differences in MNase-seq signal at the rDNA locus were more subtle than what is typically observed at single-copy loci, we sought to ensure that these differences are not due to differential MNase sensitivity of these nucleosomes. To this end, we compared the MNase-seq profiles for these nucleosomes in wild-type and *isw2∆ nhp10∆* strains using three different concentrations of MNase (Fig 3C, S3 Fig). The overall shapes of the MNase-seq profiles varied depending on MNase concentrations used. However, at any specific degree of digestion, the relative heights of nucleosomal sub-species for wild-type versus *isw2∆ nhp10∆* cells matched the patterns described above. These results confirmed that the observed shifts in nucleosome positions in mutants were not due to differential MNase sensitivity of these nucleosomes.

### Isw2 and Ino80 facilitate efficient firing of rDNA origin of replication

The prominent Isw2 peak around the rARS coupled with the shrinkage of NDRs over the rARS in chromatin remodeling factor mutants led us to ask whether origin activity is affected by these factors. To address this question, we performed two-dimensional (2D) gel electrophoresis probing activity of the rARS (Fig 4A, [3]). The Y arc of the 2D gel is comprised of restriction fragments in the process of being passively replicated, and the bubble arc of restriction fragments in which an origin of replication has actively fired. Therefore, the ratio of bubble to Y arc signals from asynchronously growing cells reflects the ratio of actively to passively replicated restriction fragments, and thus of origin efficiency. By this method, the ratio of rARS bubble to Y arc signal, and thus rARS origin efficiency, was greatest in the wild-type and slightly reduced in *isw2∆* cells. In contrast, origin efficiency was moderately reduced in *nhp10∆* cells and even more reduced in *isw2∆ nhp10∆* double mutants (Fig 4B). These results indicate that the Isw2 and Ino80 chromatin remodeling factors promote the efficient firing of the ribosomal origin of replication.

**Fig 4.**
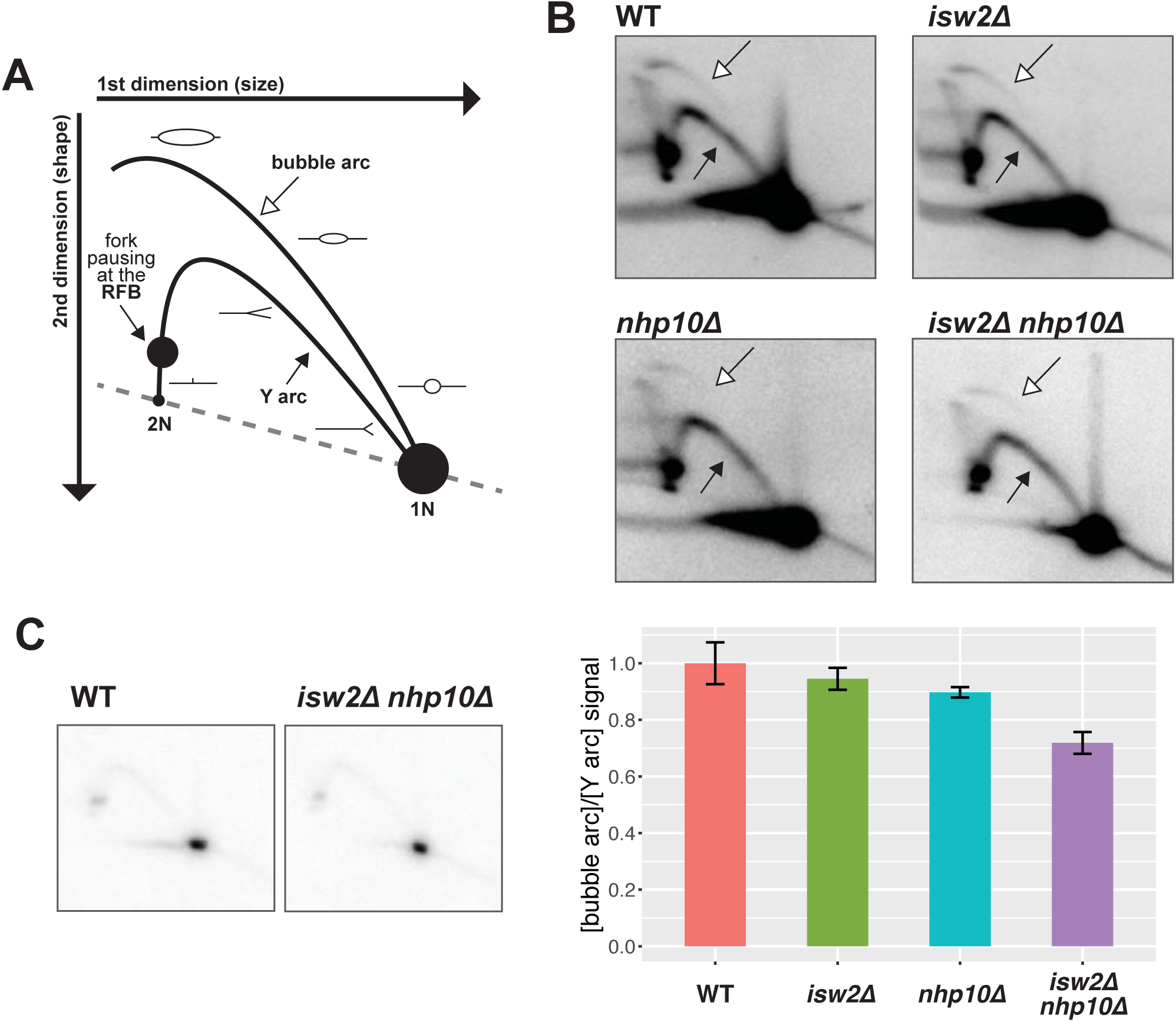
Isw2 and Ino80 facilitate efficient firing of rDNA origin of replication. (A) Schematic drawing of 2D gel with features annotated. The 1N spot is comprised of restriction fragments that are not in the process of replicating; the Y arc of restriction fragments that are being passively replicated; and the bubble arc of restriction fragments in which an origin of replication has actively fired. Replication fork pausing at the RFB causes an accumulation of restriction fragments with a specific size and shape, visible as a dark spot on the left arm of the Y-arc. The ratio of bubble arc to Y arc signal is indicative of the ratio of actively to passively replicated restriction fragments, and thus of origin efficiency. (B) Representative 2D gels over rDNA ARS and RFB. Exposures of the blots have been adjusted so that the Y arc is of comparable intensity for each blot, such that direct comparison of bubble arc intensity across images is equivalent to a comparison of bubble-to-Y ratio. Bubble arc indicated by empty arrow, Y arc indicated by filled arrow. Quantification based on measurement of average intensity of arcs using ImageQuantTL software, and reflects at least two independent experiments for each genotype. All values normalized to the bubble:Y ratio for wild-type. Error bars show standard error of the mean. (C) Representative lightly exposed 2D gel images to allow visualization of the 1N and RFB spots.

### Isw2 and Ino80 affect replication fork pausing at the rDNA Replication Fork Block

One unique aspect of DNA replication at the rDNA locus is the presence of the replication fork block (RFB). When bound by the Fob1 protein, the RFB directionally blocks the passage of replication forks from the IGS into the 3’ end of the 35S gene body, preventing head-on collisions between the replication machinery and the densely-loaded transcriptional machinery moving through the highly transcribed 35S genes [3, 7]. However, replication forks moving in the same direction as that transcriptional machinery can pass through the RFB, thus allowing for complete replication of the rDNA array. Replication fork blocking at the RFB can be detected by 2D gels as a distinct spot on the left end of the Y arc (Fig 4A). A light exposure of the 2D gel revealed reduced replication fork pausing at the RFB in remodeling factor mutants (Fig 4C). Quantifying the degree of replication fork blocking relative to the amount of loaded DNA is difficult, however, because of the large difference in the signal intensities of the RFB and the 1N spot, which represents non-replicating restriction fragments and serves as a reference for normalization. To accurately measure the degree of replication fork blocking at the RFB, we analyzed occupancy of Pol2, a subunit of DNA Polymerase epsilon, by ChIP-seq in asynchronously growing cells, an established method for globally measuring replication fork pausing [38]. Pol2 levels, and thus pausing, at the RFB are comparable to wild-type in *isw2∆* cells, but are reduced in *nhp10∆* and *isw2∆ nhp10∆* mutants (Fig 5A), similar to what we observe for rARS efficiency. In contrast, Pol2 signals at known pause sites such as *PDC1* are very similar across all tested strains (S4 Fig), suggesting that differences in pausing at the rDNA RFB are unique to that locus, and not a genome-wide phenomenon.

**Fig 5.**
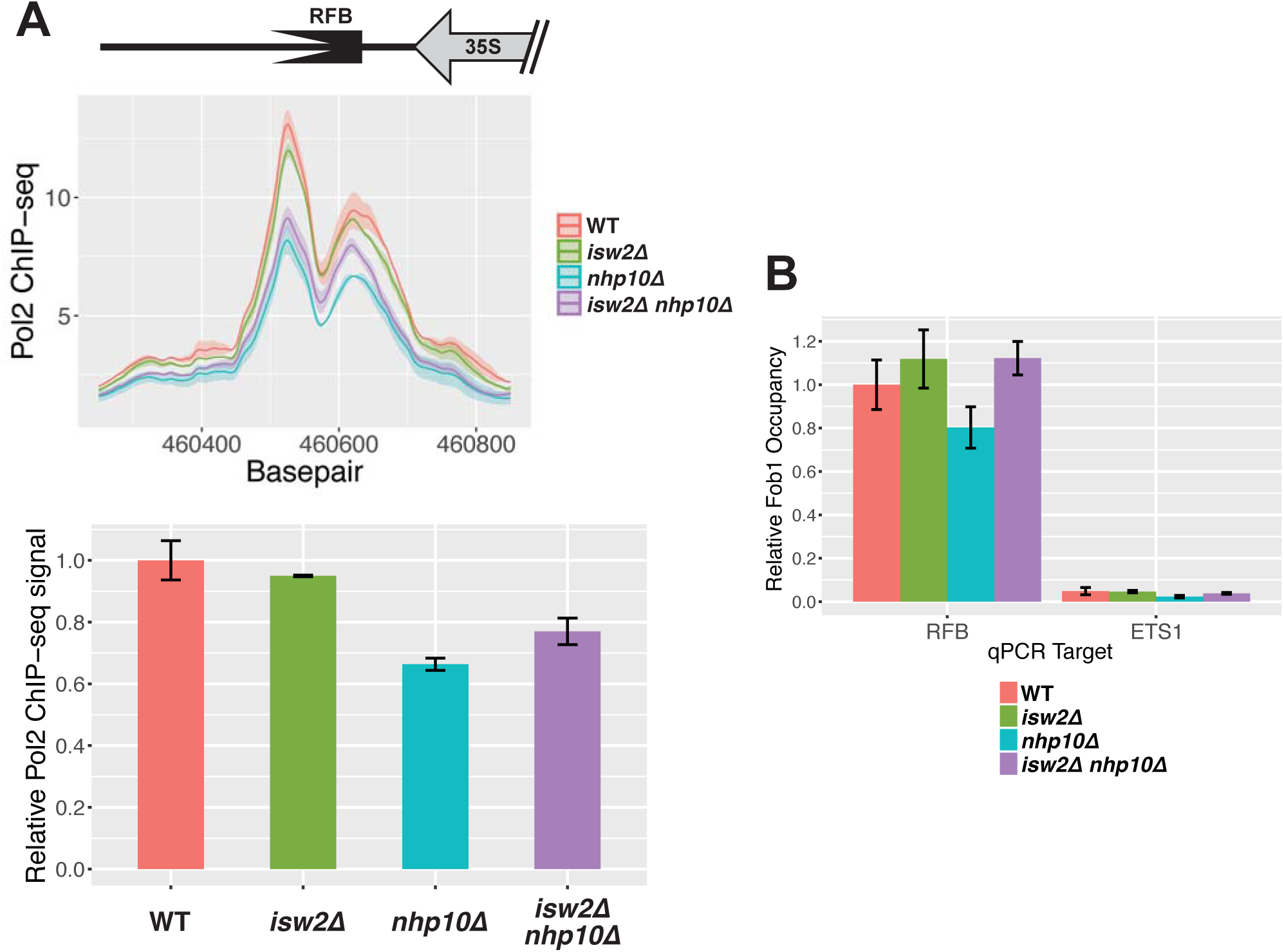
Isw2 and Ino80 affect replication fork pausing and Fob1 occupancy at the RFB. (A) ChIP-seq of DNA Polymerase Epsilon subunit Pol2 at the RFB. Ribbon plot produced as described in Fig 2A based on two biological replicates per genotype. Quantification produced by integrating the ChIP-seq signal for each strain across the RFB, and then averaging the results across two replicates. Error bars represent the standard error of the mean. (B) ChIP-qPCR of Fob1 with primers flanking the RFB and within ETS1, all normalized to occupancy at the RFB in wild-type cells. Fob1 is not expected to bind to ETS1, and thus it serves as a negative control locus. Error bars represent standard error of the mean for at least three replicates per genotype.

Because Fob1 binding at the RFB is required for pausing at this locus, we next asked whether Isw2 and Ino80 affect replication fork blocking at the RFB by altering the level of Fob1 binding. To this end, we performed ChIP of Fob1 followed by qPCR using primers flanking the RFB. This experiment revealed that the *nhp10∆* strain, which exhibits the lowest level of replication fork pausing, also shows the lowest levels of Fob1 occupancy (Fig 5B). However, *isw2∆ nhp10∆* cells, which have similarly low levels of pausing, have considerably higher levels of Fob1, on par with *isw2∆* cells and above that of the wild-type cells. Therefore, the level of Fob1 binding alone cannot explain the strain-specific differences we observe in replication fork pausing at the RFB.

### Isw2 and Ino80 affect the rate of rDNA copy number change

Replication fork pausing at the RFB is an essential step in the mechanism by which rDNA copy number is regulated. With some frequency, a targeted DNA double-strand break (DSB) will occur at replication forks paused at the RFB. Depending on how this DSB is repaired, an rDNA repeat can be removed from or added to the rDNA array, or there can be no change in rDNA copy number. Thus, Fob1-dependent replication fork pausing is a critical feature of rDNA copy number change. Given the differential pausing at the RFB in our remodeling factor mutants, we wondered whether Isw2 and Ino80 affect rDNA copy number change. To answer this question, we employed a strain in which endogenous *FOB1* has been deleted and the rDNA array reduced to 20 repeats. In the absence of Fob1, there is no pausing at the RFB, stabilizing the rDNA copy number. These cells can survive with 20 copies of the rDNA, but introduction of Fob1 via a plasmid causes rapid increase in the number of rDNA repeats via homologous recombination until the rDNA array reaches a normal size of approximately 150 copies [8]. Starting with a *fob1* strain with 20 copies of the rDNA, *ISW2*, *NHP10*, or both genes were deleted. The Fob1 gene was then reintroduced on a plasmid, and the cells were cultured continuously under selection for almost 200 generations, with samples taken at multiple time points. The copy number of rDNA repeats was monitored by CHEF gel electrophoresis followed by Southern blot analysis using a probe against the rDNA locus.

Although all four strains began to increase their rDNA copy number immediately following introduction of plasmid-borne Fob1, each of the strains behaved differently (Figs 6A, B). In wild-type and *isw2∆* cells, and to a slightly lesser degree in *nhp10∆* cells, there was a significant jump in copy number at around 35 generations after Fob1 re-introduction, the earliest time point we were able to sample. In contrast, the double mutant exhibited only a very small increase in copy number at 35 generations. After nearly 200 generations in the presence of Fob1, both the wild-type and *isw2∆* strains had recovered essentially wild-type rDNA copy number of around 150 copies, and *nhp10∆* was close to this number. In contrast, *isw2∆ nhp10∆* had barely reached 100 copies by this time point. Based on this data, we conclude that Isw2 and Ino80 facilitate the regulated increase of rDNA copy number in the rDNA array, and that their loss reduces the rate at which rDNA copy number can be increased in a population of cells.

**Fig 6.**
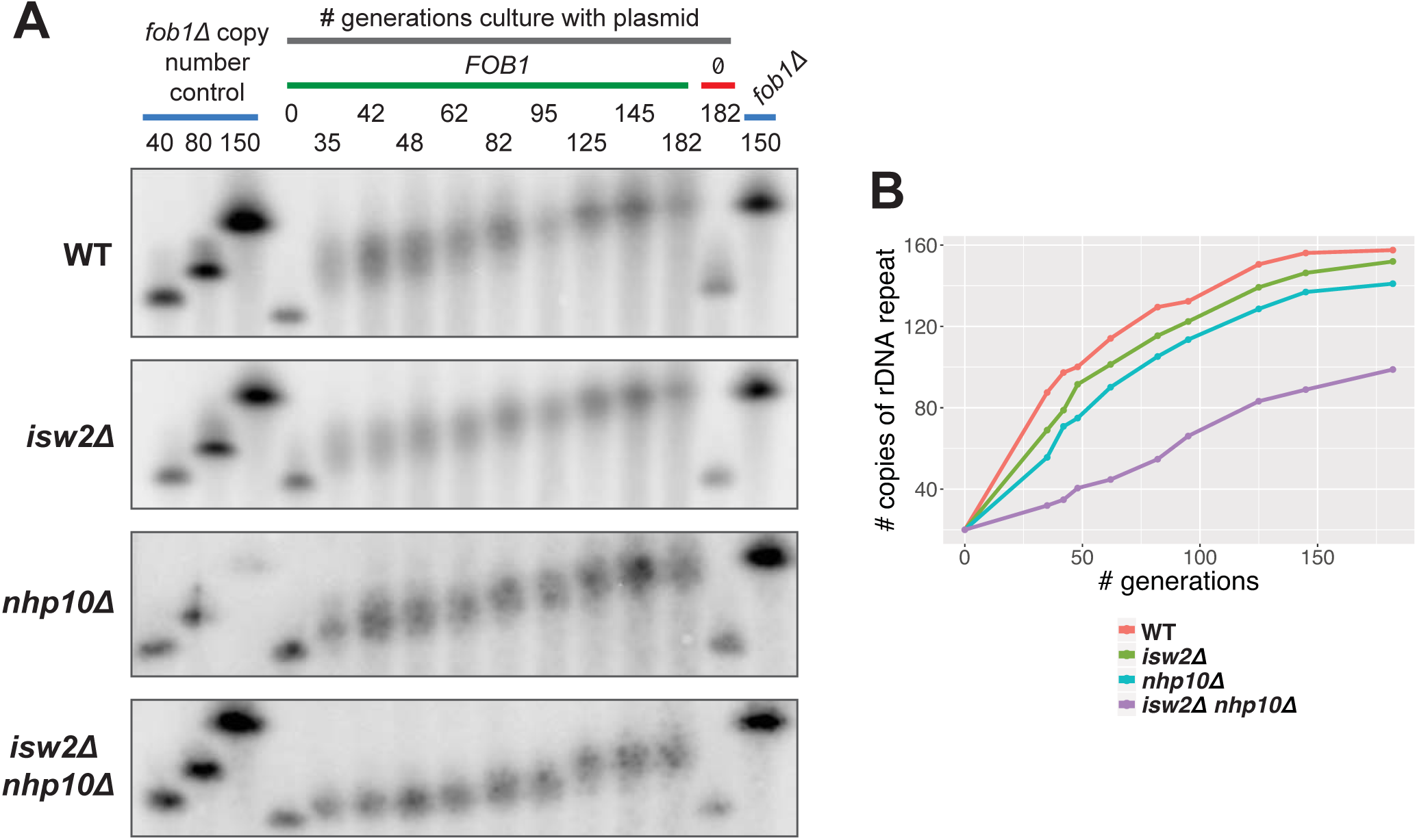
Isw2 and Ino80 affect the rate of rDNA copy number change. (A) rDNA copy number change assay. Blue bars indicate *fob1* copy number control strains that stably contain the indicated number of rDNA repeats (identical 150-copy control samples run on both ends of the gel to facilitate comparison of band migration). The gray bar denotes samples grown in a time course for the indicated number of generations, in selective medium to ensure retention of either a plasmid containing *FOB1* (green bar) or the plasmid backbone pRS426 without *FOB1* (red bar). (B) Quantification of the copy number change assay. Average copy numbers at each time point were calculated based on migration of bands relative to controls.

## Discussion

The ribosomal DNA locus is the evolutionarily conserved site of many different DNA-dependent processes, all of which must be carefully balanced. Sufficient rRNA must be transcribed to support ribosome biogenesis, but without interfering with faithful replication of the rDNA [1]. The rDNA array must be fully replicated, while still allowing for the replication of other parts of the genome [5]. The size of the rDNA array must be carefully maintained through recombination, yet the array must be protected from unintended recombination despite its highly repetitive nature. Despite many studies detailing these complex processes, relatively little is known about how ATP-dependent chromatin remodeling factors dynamically regulate chromatin structure at the *S. cerevisiae* rDNA locus to allow for these processes to occur. It has been shown that the SWI/SNF complex localizes to the rDNA and that deletion of its Snf6 subunit reduces 35S rRNA transcription [13]. In addition, it was shown that Isw2, Isw1, and Chd1 are present at the rDNA, and that their simultaneous deletion reduces 35S rRNA transcriptional termination [14]. However, the nature of chromatin regulation by these remodeling factors at the rDNA locus remains unknown, as does their involvement in processes beyond transcription of rRNA. In this study, we show that in addition to Isw2, the Ino80 ATP-dependent chromatin remodeling factor is targeted to this highly repetitive genomic locus. We show for the first time that these factors modify local chromatin structure at the levels of nucleosome occupancy, the ratio of nucleosome-occupied to nucleosome-depleted rDNA repeats, and nucleosome positioning. In addition, we find that these chromatin remodeling factors affect two critical activities that take place at the rDNA: replication initiation from the ribosomal ARS, and rDNA array amplification.

Our data indicate that Isw2 and Ino80 do not affect overall levels of 35S rRNA transcription, a result that initially surprised us. According to the prevailing model, nucleosome occupancy through the 35S gene body dictates 35S transcription, as rDNA repeats that are heavily occupied with nucleosomes are transcriptionally silent, while repeats that are depleted of nucleosomes are transcriptionally active. Thus, based on the increased nucleosome occupancy and reduced proportion of psoralen-accessible rDNA repeats observed in *isw2∆ nhp10∆* cells relative to other tested strains, we expected that 35S rRNA transcription would be correspondingly decreased in the double mutant. The lack of an effect on transcription may be explained by the robustness of 35S transcriptional regulation: when one element of this system is disrupted, another element is adjusted to maintain the desired level of transcription. For example, in a *S. cerevisiae* strain in which the rDNA array has been reduced from a normal size of ~150 copies down to ~40 copies, loading of RNA Pol I on any given active repeat is increased, such that there is no net decrease in 35S transcriptional output [30]. Similarly, in mammalian cells, inducing silencing of some rDNA repeats by depletion of UBF leads to a compensatory increase in transcription per active repeat [39]. We therefore speculate that the robust homeostatic regulation of rRNA transcription overcomes changes in nucleosome occupancy in *isw2∆ nhp10∆* cells, reacting to a reduced proportion of active repeats by increasing RNA Pol I transcription in each active unit. This would produce no net alteration in rRNA production compared to wild-type cells. We also note that our H3 ChIP-seq data reveals that the nucleosome-depleted region (NDR) in the 35S promoter is much deeper in *isw2∆ nhp10∆* cells than in wild-type or single mutant cells (S5 Fig). According to a general paradigm of RNA Pol II transcription, promoter NDR depth correlates positively with transcription [40, 41]. This deepened NDR in *isw2∆ nhp10∆* cells may reflect significantly increased loading of Pol I transcriptional machinery in each active repeat, as needed to maintain proper levels of 35S transcription despite the reduced number of active repeats.

A critical transcriptional regulator at the mammalian rDNA is the Nucleolar Remodeling Complex (NoRC), which contains SNF2h, the mammalian orthologue of yeast Isw2. Among other activities that influence rRNA transcription, this complex shifts the nucleosome at the promoter of the 45S rRNA gene, the mammalian orthologue of the yeast 35S, into a transcriptionally repressive position [12]. Notably, we see nearly identical nucleosome positioning profiles at the comparable nucleosome in *isw2∆* and *isw2∆ nhp10∆* cells compared to wild-type cells (S2D Fig). This finding, in conjunction with our observing no differences in rRNA transcription in these deletion strains, distinguishes the Isw2-mediated regulation of the yeast rDNA from the NoRC-mediated regulation of the mammalian rDNA.

While we find that loss of Isw2 and Ino80 does not affect net rRNA transcription, we do find that their loss reduces the activity of the rARS. There are multiple reports that chromatin structure around replication origins significantly affects DNA replication. Blocking an ARS with a nucleosome reduces the efficiency of that ARS [42], and proper positioning of nucleosomes adjacent to an ARS is important for replication initiation [43]. Compared to naked DNA, chromatinized DNA facilitates much greater origin selectivity at the stage of origin licensing, suggesting that chromatin structure regulates which origins fire during S-phase [44]. Consistent with these findings, ATP-dependent chromatin remodeling factors contribute to regulating replication initiation. For example, the SWI/SNF complex is targeted to a subset of origins in HeLa cells [45] and facilitates replication initiation at one out of four natural ARSs tested in a mini-chromosome maintenance assay in *S. cerevisiae* [46]. By applying an *in vitro* replication assay to nucleosomal templates remodeled by different chromatin remodeling factors, a recent study found that most factors permitted origin licensing, but that Isw2 and Chd1 prevented it [47]. As far as we know, however, there have been no reports of chromatin remodeling factors affecting both chromatin structure and replication initiation at a specific origin of replication at its natural genomic locus *in vivo*.

We report that loss of *ISW2* and *NHP10*, individually and together, reduced the efficiency of the rARS during logarithmic growth conditions in rich medium. We found that *isw2∆ nhp10∆* cells have the most robust differences in nucleosome positioning compared to wild-type cells, with a clear trend of an enrichment for nucleosomes in positions that encroach on the rARS. These same cells have the most reduced efficiency at this ARS compared to wild-type. This effect is reminiscent of the behavior of Isw2 at Pol II-transcribed genes targeted by Isw2. At such genes, when *ISW2* is deleted, NDRs at the end of the gene targeted by Isw2 tend to widen, and nearby non-coding transcription increases, suggesting that this remodeling factor typically functions to narrow these NDRs and repress non-coding transcription [18]. We observe a similar but oppositely oriented trend at the rARS, as our data suggest that the NDR containing the ARS overall becomes narrower and origin efficiency goes down in *isw2∆ nhp10∆* cells, suggesting a normal function of these factors in keeping this NDR wide and thus permissive to replication initiation. We also note that although we see intermediate effects on both nucleosome positioning and efficiency of the rARS in *isw2∆* and *nhp10∆* single mutant cells, there does not appear to be a clear additive effect that accounts for what we observe in the double mutant. Although *isw2∆* cells have more rARS-encroaching nucleosomes than *nhp10∆* cells, efficiency of the rARS appears greater in *isw2∆* cells than in *nhp10∆* cells. Thus, it appears that nucleosome positioning around the rARS can only partially account for the effect these remodeling factors have on efficiency of the rARS. Reduced rARS efficiency in our mutants may also be partially explained by the altered ratio of transcriptionally active to inactive rDNA repeats. Evidence suggests that rARSs are more likely to fire when they are adjacent to actively transcribed rDNA repeats [48]. In our proposed model, the proportion of actively transcribed repeats is reduced in *isw2∆ nhp10∆* cells, and thus a reduced proportion of rARSs in the array are adjacent to actively transcribed repeats. This may contribute to the reduced origin efficiency we observe in these mutants.

In addition to regulating rARS activity, a cell must carefully calibrate the size of the rDNA array. This highly repetitive locus must be large enough to allow for the transcription of sufficient ribosomal RNA to satisfy a cell’s demand for ribosomes; in a typical yeast cell, approximately 75 copies of the rDNA are actively transcribed to satisfy this demand [1]. However, those 75 copies of the rDNA repeat must be insufficient under some circumstances, as a typical yeast rDNA array contains around 150 copies of the rDNA repeat. According to the prevailing model, these additional copies are necessary to maximize genome stability. Active ribosomal RNA genes are transcribed at extremely high levels, with densely loaded transcriptional machinery. This presents an obstacle to the repair of damage to the underlying DNA, and persistent, un-repaired damage to the rDNA array delays complete replication of the genome and progression through S-phase [6]. Thus, to maximize genome stability, the rDNA array must be large enough to support sufficient rRNA transcription without requiring all repeats to be actively transcribed. This requirement imposes a lower limit on the optimal size of the rDNA array. Similarly, the array cannot exceed a certain size. If the rDNA grows too large, its complete replication would require an excessively large proportion of the finite pool of replisome components available during each S-phase, depriving other parts of the genome of those replication factors [5]. In addition, evidence suggests that having a smaller rDNA array improves growth during persistent replication stress, perhaps by making more of the limiting replication factors available to other parts of the genome [49]. Thus, the number of repeats in the rDNA locus must be actively managed by the cell to facilitate optimal transcriptional output and maximize genome stability. Most of our knowledge about the mechanism of rDNA copy number change comes from studying the cellular response to a significant perturbation in copy number. For example, if an rDNA array is artificially truncated, it will steadily increase until it reaches a normal size [50]. Conversely, the rDNA array will shrink when the *RPA135* subunit of RNA Pol I is deleted [50, 51], when the activity of the origin recognition complex is compromised [52], or when a number of other replication factors are lost [49]. Together, these studies demonstrate that maintenance of the size of the rDNA is a vital process that is actively regulated by the cell.

In this study, we describe a nearly two-fold reduction in the rate of copy number increase in *isw2∆ nhp10∆* cells relative to wild-type cells, and moderate reductions in the rate of increase in *isw2∆* and *nhp10∆* cells. This is the first demonstration of any ATP-dependent chromatin remodeling factors contributing to the regulation of rDNA copy number change. We show that these remodeling factors affect Fob1 binding and replication fork pausing at the RFB, two critical steps in the process of copy number change, but the effects on these activities do not clearly correlate with the effects on the rate of copy number change we observe in the same mutants. Therefore, we do not believe the remodeling factors influence copy number change exclusively through replication fork pausing or Fob1 binding. Another critical step in this process is the repair of the targeted DNA double strand break (DSB) that takes place at RFB-paused replication forks. In light of a well-documented role for Ino80 in DSB repair [21, 53, 54], it is possible that the striking defect in copy number increase we observe may result in part from mis-regulation of the recombination-based repair of these DSBs. In the absence of *NHP10*, there may be some mild defect in homologous recombination (HR) that may be partially compensated for by otherwise normal chromatin structure created by Isw2. However, in the double mutant, HR repair defects may become too significant to facilitate the desired recombination rate at rDNA.

In addition to this possible direct involvement of the remodeling factors in copy number increase, the effect we observe may be indirect. Our data suggest that in the absence of Isw2 and Ino80, the ratio of active to inactive rDNA repeats is reduced. In light of work showing that rARSs are more likely to fire when they are adjacent to actively transcribed rDNA repeats [48], we proposed above that this reduced proportion of active repeats could explain the reduced efficiency of the rARS in *isw2∆ nhp10∆* cells. It has also been shown that copy number change events require firing of the rARS adjacent to the RFB at which a replication fork is paused, a DSB is induced, and then repaired. This same study found that the efficiency of the ARS in the IGS correlates with the rate of copy number increase [55]. Accordingly, it is possible that the reduced ratio of active to inactive repeats in the double mutant causes a change in rARS efficiency, which in turn reduces the frequency of copy number change events, thus accounting for the reduced rate of copy number increase in the double mutant cells. In sum, this work establishes a novel role for ATP-dependent chromatin remodeling factors in strongly influencing rDNA biology, including the process of rDNA copy number change.

## Materials and Methods

### Yeast strains and media

Strains used are listed in S1 Table. Strains generated using standard gene replacement protocols. Unless otherwise indicated, yeast cells were grown in YEPD medium (2% Bacto Peptone, 1% yeast extract, 2% glucose). All strains *MAT***a** W303-1a.

### Chromatin immunoprecipitation and micrococcal nuclease digestion followed by deep sequencing

Chromatin immunoprecipitation (ChIP) and micrococcal nuclease (MNase) digestion were performed as described previously [58]. For H3-ChIP experiments, anti-H3 C-term antibody (Abcam catalog # ab1791) was used; for all other ChIPs, the targeted protein was epitope-tagged with FLAG, and immuno-precipitated using anti-FLAG monoclonal antibody (Sigma catalog # F3165). All Isw2 ChIP-seq performed on a FLAG-tagged, catalytically inactive allele of *ISW2* as previously described [59]. All libraries were constructed using the Nugen Ovation Ultralow System V2 (catalog # 0344-32) and then subjected to single-end (ChIP-seq) or paired-end (MNase-seq) sequencing, with 50 bp read length, on Illumina Hi-Seq 2500. Ribbon plots, bar graphs, and line graphs were generated with the ggplot2 R package (http://ggplot2.org/). For all depictions of deep-sequencing data at the rDNA, a single copy of the rDNA locus is shown. Our reference genome contains two copies of the rDNA, and any read mapping to the rDNA is randomly assigned to one of these 2 copies. Thus, sequencing data reflects the average signal across all rDNA repeats in all cells sampled.

### Reverse Transcription- and ChIP-quantitative PCR

RNA was isolated using hot acid phenol, then cleaned up with the Qiagen RNEasy Cleanup Kit (catalog # 74204) plus on-column treatment with DNase I (Qiagen catalog # 79254). cDNA was generated from the RNA using Superscript III Reverse Transcriptase (ThermoFisher catalog # 18080093). Quantitative PCR was performed on both cDNA and ChIP DNA using 2x Power SYBR Master Mix (Fisher Scientific catalog # 4367659) run on the ABI QuantStudio5 Real Time PCR System machine.

### Psoralen Crosslinking

Assay was performed as previously described [10, 28, 32]. Cells were grown to mid-log phase (OD_660_ = 0.5-0.7), approximately 3×10^8^ cells were collected, washed twice with ice cold water, and then re-suspended in 1.4 ml cold TE buffer. Cells were transferred to 6 well plates, and 70 ul of psoralen (200 ug/ml in 100% ethanol) was added to the cells. On ice, the plates were irradiated with 365 nm UV for five minutes. Psoralen addition followed by UV irradiation was repeated four additional times, for a total of five rounds. Cells were collected, washed in water, spheroplasted with zymoylase 100T, and washed in spheroplast buffer. The pellet was lysed by re-suspension in TE buffer with 0.5% SDS and then treated with Proteinase K overnight at 50°C. DNA was extracted with Phenol:Chloroform:IAA, ethanol precipitated, and then digested for at least 3 hours with EcoRI-HF. Samples were treated with RNase A at 37°C for 30 minutes, ethanol precipitated, quantified, and then run in 1.3% LE agarose gels in 0.5X TBE for 24 hours at 60V. Gels were irradiated for two minutes per side with a Stratagene Stratalinker, transferred to a GeneScreen Plus membrane in 10x SSC, and then hybridized with a probe contained within a EcoRI restriction fragment in the rDNA ETS1. Membranes were visualized using a Typhoon Phosphor Imager, and images were visualized using ImageJ software.

### 2D gel electrophoresis

DNA sample preparation based on the Brewer/Raghuraman lab protocol (http://fangman-brewer.genetics.washington.edu/plug.html). Cells were grown to mid-log phase (OD_660_ = 0.5-0.7), sodium azide added to 0.1% final concentration, and then cultures were washed in water. Cell pellets were re-suspended in 50 mM EDTA, mixed with an equal volume of 1.0% Low-Melt Agarose (BioRad catalog # 161-3111), and pipetted into plug molds. Cells in plugs were spheroplasted with 0.5 mg/ml Zymolyase 20-T, thoroughly washed, and stored in TE at 4°C. Plugs were digested with NheI for 5 hours at 37°C, then run in 0.4% agarose gels in TBE at 1 V/cm for 22 hours at room temperature. Gels were stained with ethidium bromide (EtBr), visualized with UV, and the desired size range for each sample was identified in the gel and physically cut out. This piece of gel was then rotated 90° and placed in a new gel tray, and warm 1.1% agarose in TBE was poured around it. This gel was then run at 5 V/cm for 6 hours at 4°C. After running, the gel was visualized, transferred onto a GeneScreen Plus membrane (Perkin Elmer, catalog # NEF986001PK), and hybridized with a probe encompassing the RFB.

### rDNA copy number change assay

Strains were made from YSI102 [6], in which the endogenous *FOB1* gene had been deleted, and the number of rDNA repeats reduced to 20 copies. From the 20-rDNA-copy *fob1* parent, *isw2∆*, *nhp10∆*, and *isw2∆ nhp10∆* strains were generated. Separately, the *FOB1* gene was cloned into the pRS426 plasmid using Gibson cloning. Either this *FOB1*-pRS426 plasmid or a pRS426 plasmid with no *FOB1* gene was then transformed into each 20-copy strain and plated on yeast complete (YC) –URA medium with 2% glucose. Individual transformants were re-streaked on selective medium, presence of the desired plasmid was confirmed by PCR, and then transformants were inoculated into liquid YC –URA + 2% glucose. Cultures were allowed to reach saturation, and then aliquots were collected, washed in cold 50 mM EDTA, and cell pellets were frozen in liquid nitrogen and stored at −80°C. From the remaining saturated cultures, all strains were diluted by the same factor, then allowed to grow back to saturation, at which point the next time point would be collected, up to ~200 generations. Generations were calculated from the base 2 log of the dilution factor applied at each passage (e.g. a saturated culture diluted by a factor of 1,024 into the same volume of medium would require 10 generations to return to saturation).

### Clamped Homogenous Electric Field (CHEF) gel electrophoresis and Southern blotting

Samples for CHEF gels were prepared in agarose based on a previously described method [60]. Frozen cell pellets were thawed in room-temperature water, re-suspended in 100 mM EDTA, then mixed with 0.8% Low-Melt Agarose and 25 mg/ml zymolyase 20T. This mixture was pipetted into plug molds, allowed to solidify at 4°C, then washed with a series of buffers (Solution V: 500 mM EDTA pH 7.5, 10 mM Tris pH 7.5; Solution VI: 5% sarcosyl, 5 mg/ml proteinase K, 500 mM EDTA pH 7.5; Solution VII: 2 mM Tris pH 7.5, 1 mM EDTA, pH 8.0). Before being run, plugs were incubated for approximately 30 minutes in TBE running buffer at 4°C before being placed on gel comb teeth, positioned in gel mold, and then warm 0.8% 0.5x TBE was poured. CHEF gel was run on a CHEF-DR II with a program adapted from Ide *et al* MCB 2007: Block 1 = 2.0 V/cm, pulse time of 1,200 seconds to 1,400 seconds, total run time 72 hours; Block 2 = 6.0 V/cm, pulse time of 25 seconds to 146 seconds, total run time 7.5 hours. After electrophoresis, gels were incubated with 0.5 ug/ml EtBr in running buffer for 30-45 minutes, UV-irradiated with a Stratagene Stratalinker to nick DNA, transferred onto HyBond N+ positively charged membrane (GE, catalog # RPN303B), and hybridized with a probe targeting the RFB.

## Acknowledgments

We are grateful to members of the Brewer/Raghuraman lab, especially Bonny Brewer, MK Raghuraman, Joe Sanchez, and Liz Kwan, for help with experimental design and troubleshooting, as well as conceptual discussion and feedback, and members of the Smith lab, especially Randy Hyppa, for help with CHEF gels. We thank Takehiko Kobayashi for strains used in this study. Finally, we thank members of the Tsukiyama lab for helpful discussion and commentary on this manuscript.

## Supporting Information

**S1 Fig.**
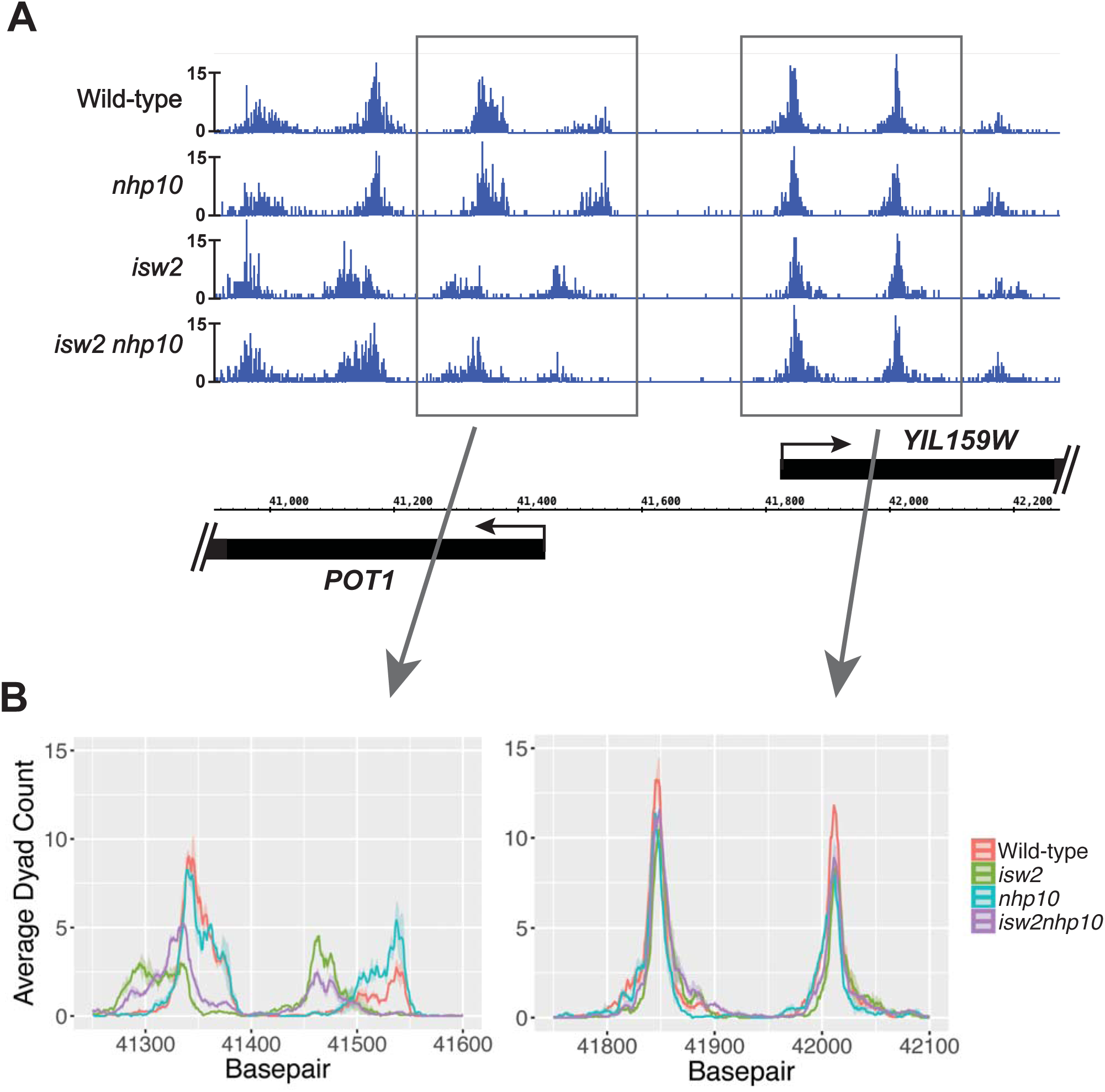
Striking nucleosome positioning changes at a canonical Isw2 target. (A) MNase-seq data at a well-established Isw2 target, the 5’ end of the *POT1* gene. (B) Visualization of the same data shown in S2A Fig with the ribbon plots used in Fig 3B, focusing on two pairs of nucleosomes.

**S2 Fig.**
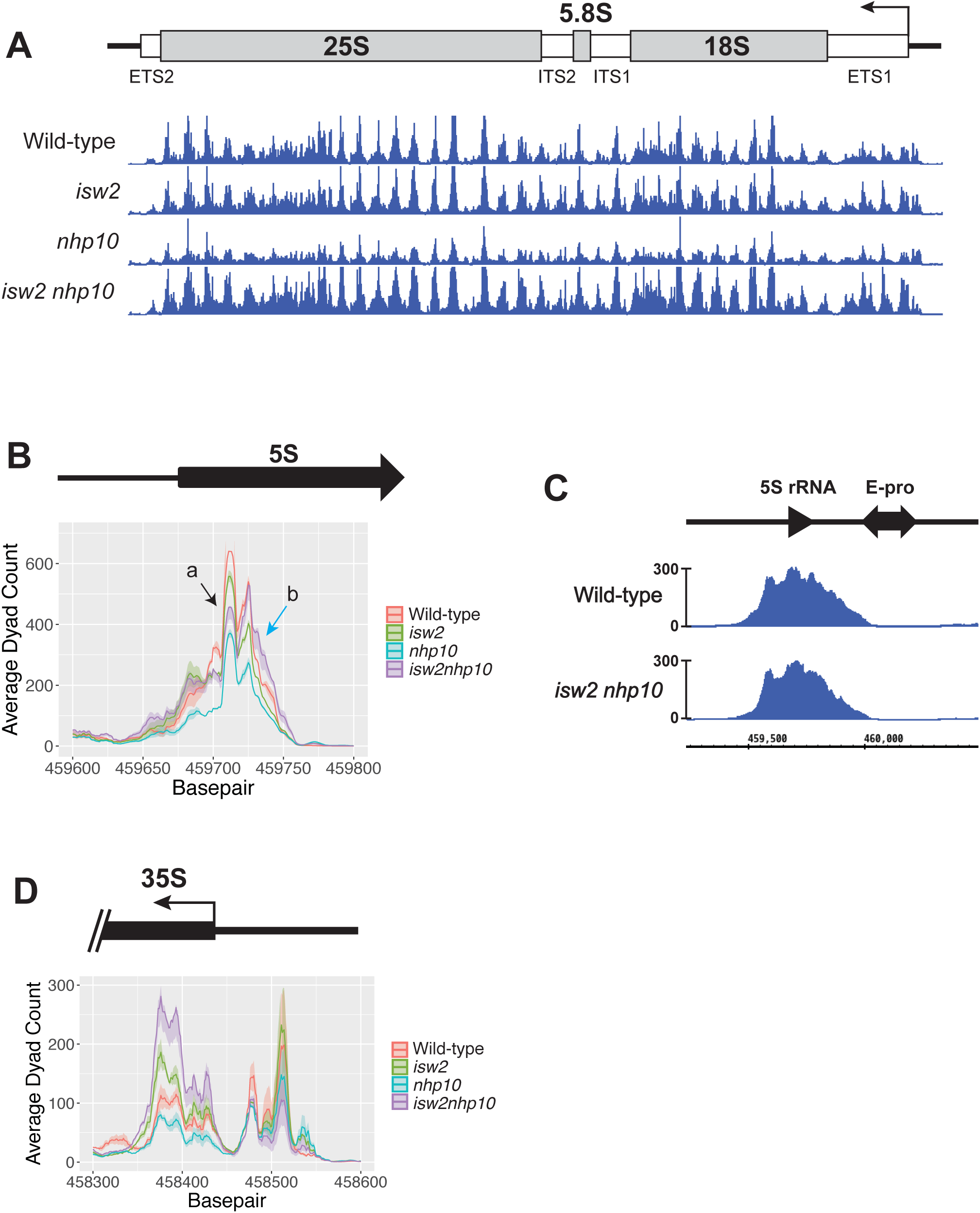
Nucleosome positioning in single and double mutants throughout the rDNA. (A) MNase-seq data analyzed with dyad mapping showing the entire 35S rRNA gene. No clear differences in nucleosome positioning can be seen. (B) Ribbon plot of the same MNase-seq data focused on the 5S-adjacent nucleosome. In wild-type cells and *isw2∆* and *nhp10∆* cells, sub-species *a* is higher than sub-species *b*, while in *isw2∆ nhp10∆* cells the *b* peak is higher than for *a*. (C) RNA Pol III ChIP-seq at the Pol III-transcribed 5S gene showing no appreciable difference in levels of the polymerase between wild-type and *isw2∆ nhp10∆* cells. (D) MNase-seq ribbon plot at the 35S promoter region showing no appreciable difference in nucleosome positioning across the strains tested.

**S3 Fig.**
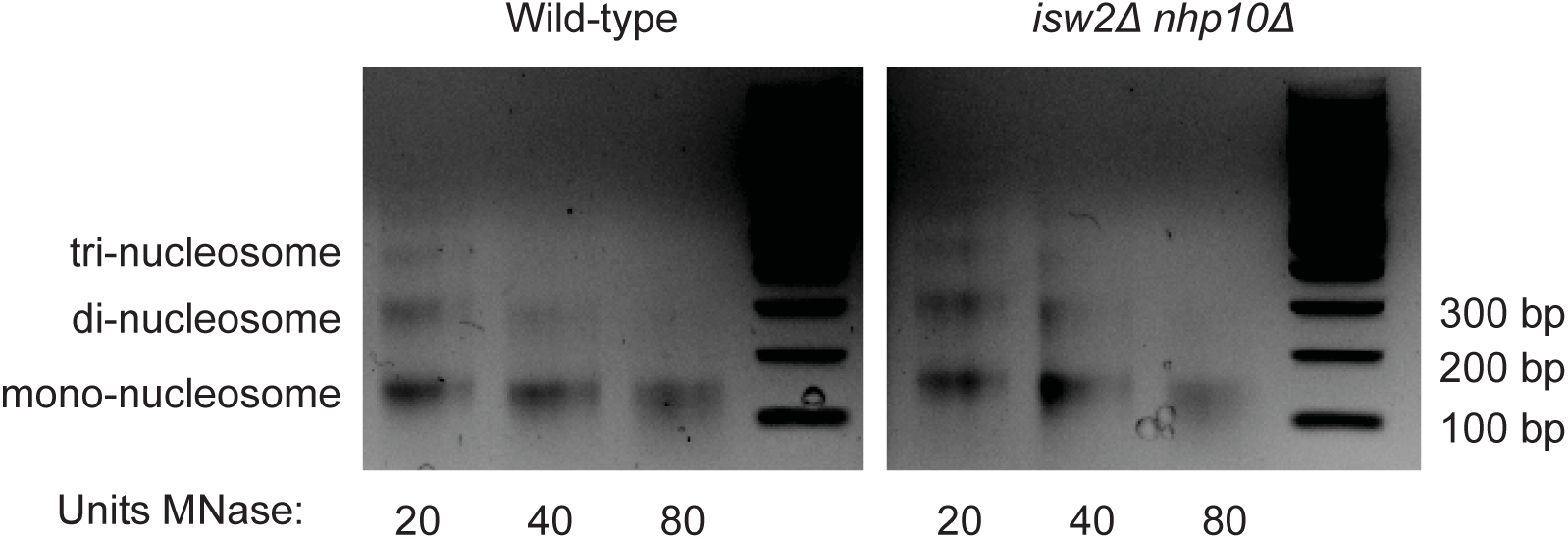
Different MNase digestions. (D) Representative gel indicating how nucleosomal ladders appear after digestion with 20, 40, or 80 units of MNase. Note that for all MNase-seq analyses, regardless of level of digestion, the mono-nucleosomal band was gel-purified and was the sole source of material subjected to deep sequencing.

**S4 Fig.**
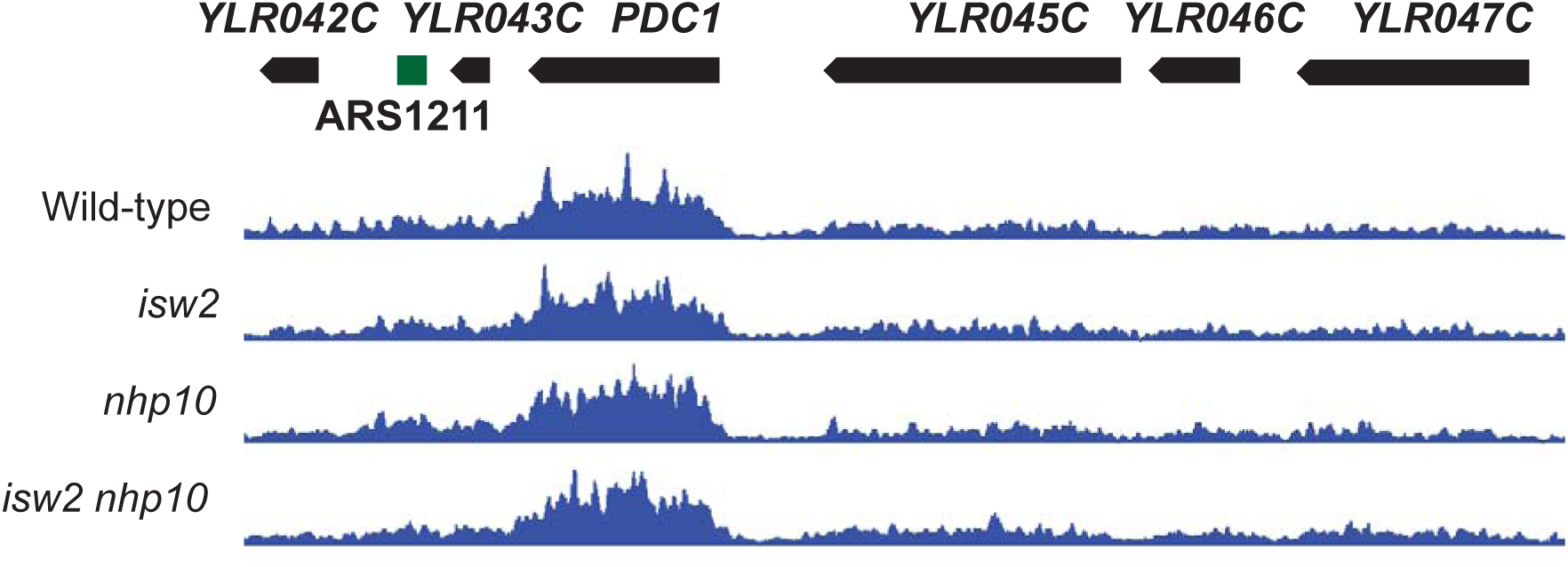
DNA polymerase pausing does not vary between tested strains at a known pause site, *PDC1*. ChIP-seq of DNA polymerase epsilon subunit Pol2 at the *PDC1* gene, a site known to show polymerase pausing by this method (Azvolinsky 2009). Levels of Pol2 do not appreciably vary across the strains tested.

**S5 Fig.**
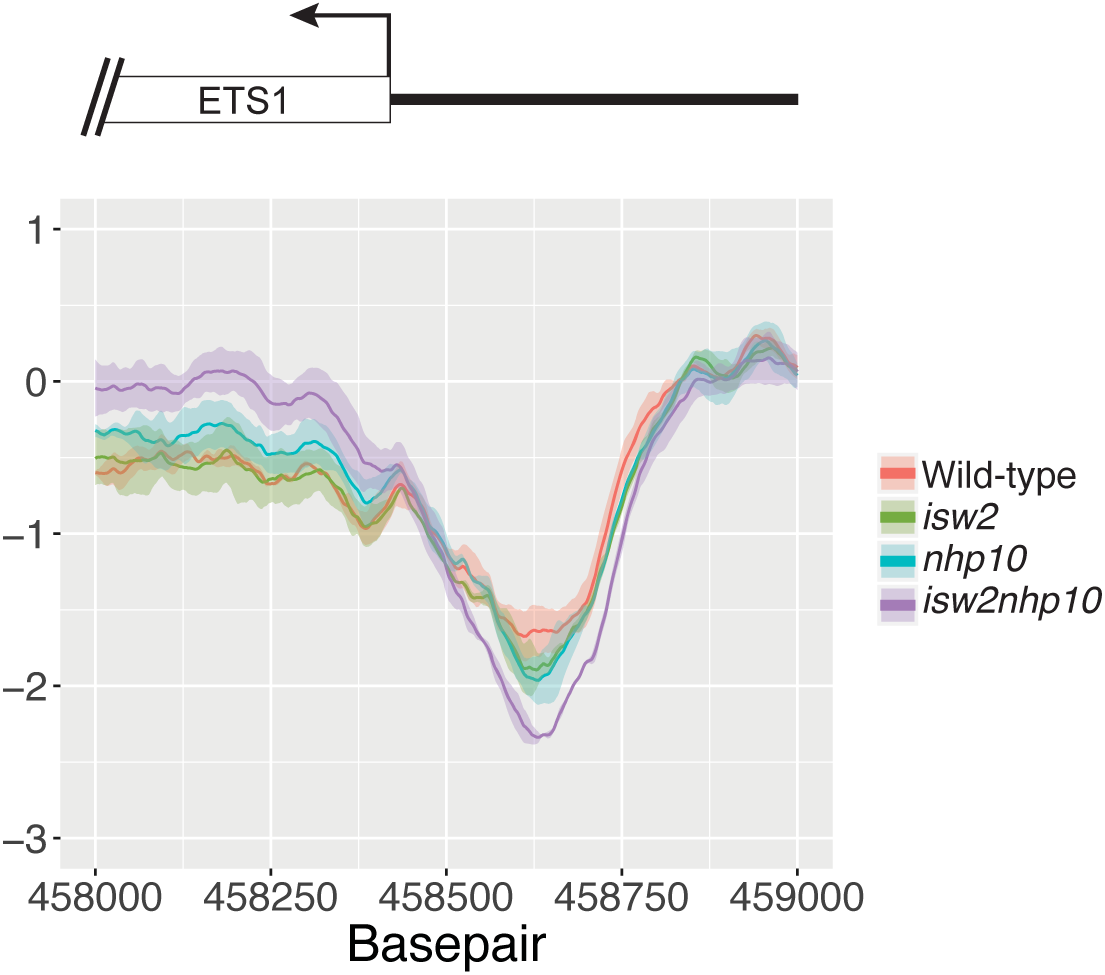
Isw2 and Ino80 affect depth of the 35S promoter’s nucleosome depleted region. The same H3 ChIP-seq data shown in Fig 2A, zoomed in on the 35S promoter region (Y axis on log2 scale). Depth of the NDR is appreciably greater in *isw2∆ nhp10∆* cells than in wild-type cells, and NDR depth is at an intermediate level in single mutant cells.

**S1 Table.** List of yeast strains used.

| Strain | Genotype | Reference |
| --- | --- | --- |
| W1588-4C | *MATa ade2-1 can1-100 his3-11,15 leu2-3,112 trp1-1 ura3-1* | Thomas and Rothstein 1989, Zhao *et al* 1998 |
| YTT3320 | W1588-4C; *isw2Δ::NatMX* | Au *et al* 2011 |
| YTT6809 | W1588-4C; *isw2Δ::NatMX* | this study |
| YTT3333 | W1588-4C; *nhp10Δ::Hyg* | Au *et al* 2011 |
| YTT2060 | W1588-4C; *nhp10Δ::Hyg* | Vincent *et al* 2008 |
| YTT3337 | W1588-4C; *isw2Δ::NatMX nhp10Δ::HYG* | Au *et al* 2011 |
| YTT2109 | W1588-4C; *isw2Δ::NatMX nhp10Δ::HYG* | Vincent *et al* 2008 |
| YTT1996 | W1588-4C; *ISW2-K215R-3FLAG-KanMX* | Gelbart *et al* 2005 |
| YTT1997 | W1588-4C; *ISW2-K215R-3FLAG-KanMX* | Gelbart *et al* 2005 |
| YTT3426 | W1588-4C; *NHP10-3FLAG-KanMX* | Vincent *et al* 2008 |
| YTT3427 | W1588-4C; *NHP10-3FLAG-KanMX* | Vincent *et al* 2008 |
| YTT6639 | W1588-4C; *rpa49Δ::KanMX* | this study |
| YTT6673 | W1588-4C; *isw2Δ::NatMX nhp10Δ::Hyg RPA190-2L-3FLAG::KanMX* | this study |
| YTT6679 | W1588-4C; *RPA190-2L-3FLAG::KanMX* | this study |
| YTT6686 | W1588-4C; *RPO31-2L-3FLAG::KanMX* | this study |
| YTT6693 | W1588-4C; *isw2Δ::NatMX nhp10Δ::Hyg RPO31-2L-3FLAG::KanMX* | this study |
| YTT6915 | W1588-4C; *Pol2-2L-3FLAG::KanMX* | this study |
| YTT6916 | W1588-4C; *Pol2-2L-3FLAG::KanMX* | this study |
| YTT6917 | W1588-4C; *isw2Δ::NatMX Pol2-2L-3FLAG::KanMX* | this study |
| YTT6918 | W1588-4C; *isw2Δ::NatMX Pol2-2L-3FLAG::KanMX* | this study |
| YTT6919 | W1588-4C; *nhp10Δ::Hyg Pol2-2L-3FLAG::KanMX* | this study |
| YTT6920 | W1588-4C; *nhp10Δ::Hyg Pol2-2L-3FLAG::KanMX* | this study |
| YTT6921 | W1588-4C; *isw2Δ::NatMX nhp10Δ::Hyg Pol2-2L-3FLAG::KanMX* | this study |
| YTT6922 | W1588-4C; *isw2Δ::NatMX nhp10Δ::Hyg Pol2-2L-3FLAG::KanMX* | this study |
| YTT7009 | W1588-4C; *Fob1-2L-3FLAG::KanMX* | this study |
| YTT7010 | W1588-4C; *Fob1-2L-3FLAG::KanMX* | this study |
| YTT7011 | W1588-4C; *isw2Δ::NatMX Fob1-2L-3FLAG::KanMX* | this study |
| YTT7012 | W1588-4C; *isw2Δ::NatMX Fob1-2L-3FLAG::KanMX* | this study |
| YTT7013 | W1588-4C; *nhp10Δ::Hyg Fob1-2L-3FLAG::KanMX* | this study |
| YTT7014 | W1588-4C; *nhp10Δ::Hyg Fob1-2L-3FLAG::KanMX* | this study |
| YTT7015 | W1588-4C; *isw2Δ::NatMX nhp10Δ::Hyg Fob1-2L-3FLAG::KanMX* | this study |
| YTT7016 | W1588-4C; *isw2Δ::NatMX nhp10Δ::Hyg Fob1-2L-3FLAG::KanMX* | this study |
| YSI101 | *MATa ade2-1 can1-100 his3-11,15 leu2-3,112 trp1-1 ura3-1 fob1::LEU2* | Ide *et al* 2010 |
| YSI102 | YSI101; 20 copies rDNA | Ide *et al* 2010 |
| YSI103 | YSI101; 40 copies rDNA | Ide *et al* 2010 |
| YSI104 | YSI101; 80 copies rDNA | Ide *et al* 2010 |
| YTT6294 | YSI102; *isw2Δ::NatMX* | this study |
| YTT6865 | YSI102; *nhp10Δ::HYG* | this study |
| YTT6311 | YSI102; *isw2Δ::NatMX nhp10Δ::Hyg* | this study |
| YTT6312 | YSI102; *isw2Δ::NatMX nhp10Δ::Hyg* | this study |

**S2 Table.**
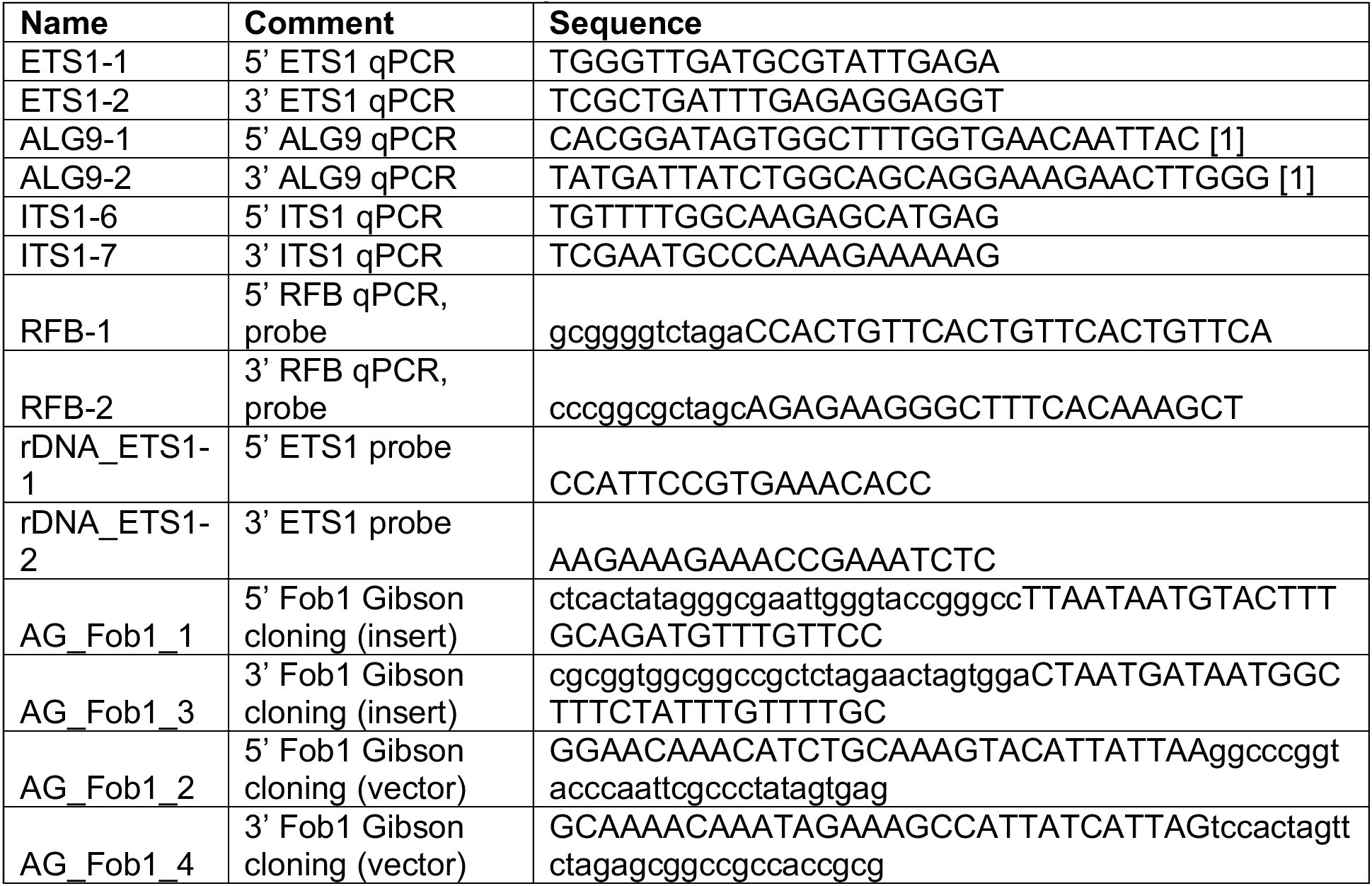
List of PCR primers used.

**S3 Table.**
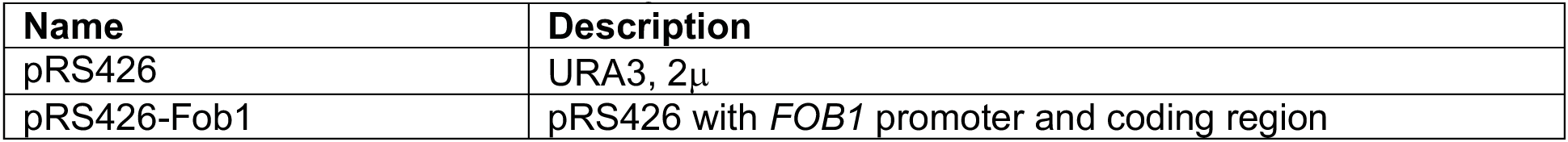
List of plasmids used.

| Name | Description |
| --- | --- |
| pRS426 | URA3, 2μ |
| pRS426-Fob1 | pRS426 with *FOB1* promoter and coding region |

